# Identification and characterization of thiamine analogues with antiplasmodial activity

**DOI:** 10.1101/2024.07.18.604204

**Authors:** Imam Fathoni, Terence C. S. Ho, Alex H. Y. Chan, Finian J. Leeper, Kai Matuschewski, Kevin J. Saliba

## Abstract

Thiamine is metabolized into thiamine pyrophosphate (TPP), an essential enzyme cofactor. Previous work has shown that oxythiamine, a thiamine analogue, is metabolized by thiamine pyrophosphokinase (TPK) into oxythiamine pyrophosphate (OxPP) within the malaria parasite *Plasmodium falciparum*, and then inhibits TPP-dependent enzymes, killing the parasite *in vitro* and *in vivo*. To identify a more potent antiplasmodial thiamine analogue, 11 commercially available compounds were tested against *P. falciparum* and *P. knowlesi*. Five active compounds were identified, but only N3-pyridyl thiamine (N3PT), a potent transketolase inhibitor and candidate anticancer lead compound, was found to suppress *P. falciparum* proliferation with an IC_50_ value 10-fold lower than that of oxythiamine. N3PT was active against *P. knowlesi* and was >17 times less toxic to human fibroblast, as compared to oxythiamine. Increasing the extracellular thiamine concentration reduced the antiplasmodial activity of N3PT, consistent with N3PT competing with thiamine/TPP. A transgenic *P. falciparum* line overexpressing TPK was found to be hypersensitized to N3PT. Docking studies showed an almost identical binding mode in TPK between thiamine and N3PT. Furthermore, we show that [^3^H]thiamine accumulation, resulting from a combination of transport and metabolism, in isolated parasites is reduced by N3PT. Treatment of *P. berghei*-infected mice with 200 mg/kg/day N3PT reduced their parasitemia, prolonged their time to malaria symptoms, and appeared to be non-toxic to mice. Collectively, our studies are consistent with N3PT competing with thiamine for TPK binding and inhibiting parasite proliferation by reducing TPP production, as well as being converted into a TPP antimetabolite that inhibits TPP-dependent enzymes.

## Introduction

Malaria is a significant disease in Africa, several countries in Asia, and in Central and South America. In 2022, malaria infected approximately 249 million people and caused 608,000 deaths (1). The spread of antimalarial drug resistance has been rapid and extensive (2, 3). The limited availability of antimalarials has prompted researchers to develop new treatments (4, 5). During its intra-erythrocytic asexual replication, which is exclusively responsible for the clinical symptoms, *Plasmodium* requires various vitamins as cofactors for biochemical processes (6). While nutrients such as amino acids and glucose are obtained from the host plasma or erythrocyte (7), thiamine **1** (vitamin B_1_) can either be synthesized by the parasite itself (8) or acquired from the host. Thiamine **1**, being positively charged, is transported into the cytoplasm, where it is converted into the cofactor thiamine pyrophosphate (TPP) **2** by thiamine pyrophosphokinase (TPK; PF3D7_0924300) (Figure 1). The important role of *Pf*TPK is highlighted by a “likely essential” score in a genome-wide piggyBac transposon mutagenesis screen (9). TPP **2** functions as an essential cofactor for multiple TPP-dependent enzymes, including pyruvate dehydrogenase E1 subunit (EC: 1.2.4.1; PF3D7_1124500; PF3D7_1446400) and 1-deoxy-D-xylulose 5-phosphate synthase (EC: 2.2.1.7; PF3D7_1337200) both located in the apicoplast, oxoglutarate dehydrogenase E1 subunit/alpha-ketoglutarate dehydrogenase (EC: 1.2.4.2; PF3D7_0820700) and branched-chain alpha ketoacid dehydrogenase E1a subunit (EC: 1.2.4.4; PF3D7_1312600) both located in the mitochondrion, and transketolase (EC: 2.2.1.1; PF3D7 0610800) located in the cytoplasm (10–15). Due to the important roles of these enzymes in cellular metabolism, the thiamine/TPP utilization pathway is a promising antimalarial target (16).

**Figure 1.**
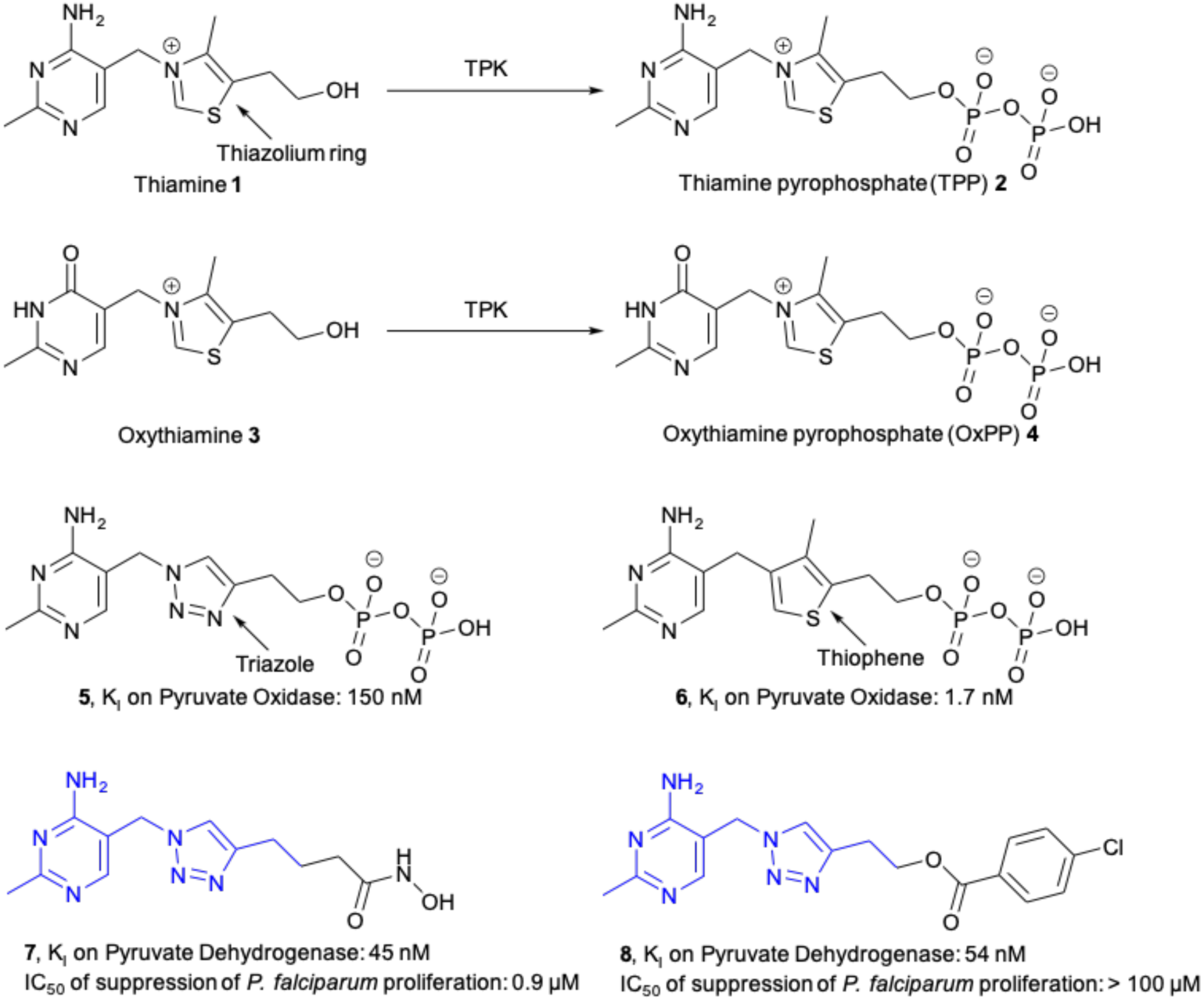
Structures of thiamine, TPP and their analogues. Key moieties, the thiazolium, triazole and thiophene rings, are indicated by the arrows. Activity of compounds **5** and **6** against pyruvate oxidase and compounds **7** and **8** against pyruvate dehydrogenase are shown. The previously-reported antiplasmodial activity of compounds **7** and **8** is also included. The common sub-structure between compounds **7** and **8** is highlighted in blue.

Several thiamine analogues with potent antiplasmodial activity have been reported. Oxythiamine **3** is a thiamine analogue, where the amino group on the pyrimidine ring is replaced by a hydroxyl group (Figure 1). Oxythiamine possesses antiplasmodial activity, both *in vitro* and *in vivo*, and has been used to validate the thiamine utilization pathway as an antimalarial drug target (15). As with **1**, **3** is activated by TPK into its pyrophosphate form, oxythiamine pyrophosphate (OxPP; Figure 1) **4**. Occupying the cofactor pocket (with its PP moiety) but lacking catalytic activity (due to its modified pyrimidine ring), **4** is an inhibitor of TPP-dependent enzymes (17). Previous work has shown that at least two TPP-dependent enzymes may be inhibited by oxythiamine **3** (after conversion to OxPP **4**) as part of its mechanism of action against *P. falciparum* parasites (15).

Replacing the positively charged thiazolium ring of TPP with a neutral ring abolishes the catalytic capability, and we have shown that the resultant TPP analogues are potent inhibitors of TPP-dependent enzymes (18–20). While TPP analogues **5** and **6** are structurally similar (Figure 1), changing the central ring from triazole to thiophene can lead to >80-fold increase in enzyme inhibition (18). Thiamine analogues **7** and **8,** bearing a common substructure (as highlighted in blue in Figure 1), are both potent enzyme inhibitors; however, only **7**, featuring a hydroxamate tail group, inhibited *P. falciparum* parasite proliferation (19, 20). Our recent findings collectively suggest that despite being structurally similar, various analogues of **1** and **2** can have disparate biological activities, highlighting the importance of even subtle changes. Therefore, in this study, we investigated the antiplasmodial activity of a range of thiamine analogues in the hope of discovering an on-target thiamine analogue with improved properties compared to **3**.

## Materials and Methods

The following thiamine analogs were purchased from Toronto Research Chemicals Canada or Sigma Aldrich: fursultiamine; cycotiamine; dibenzoyl thiamine; beclotiamine; oxothiamine; thiotiamine; bisbentiamine. N3-pyridyl thiamine (N3PT) was purchased from MedChemExpress. Allithiamine and acetiamine were kindly provided by the Developmental Therapeutic Programs (DTP) Cancer of the USA.

### Parasite culture

The human malaria parasite *P. falciparum* strain 3D7 (chloroquine-sensitive) and the same strain expressing an extra copy of TPK with a GFP-tag (*Pf*TPK-GFP) were maintained in the intraerythrocytic stage essentially as described previously (21). The macaque malaria parasite *P. knowlesi* strain H1 (adapted to human serum) was also maintained in the same way as *P. falciparum*, except that the culture medium was supplemented with 10% heat-inactivated, pooled human serum (22). Fresh human erythrocytes (blood type O^+^) were added when parasites were in the trophozoite stage. The parasite cultures were maintained at 37°C inside a shaking incubator and under an atmosphere of 1% oxygen, 3% carbon dioxide and 96% nitrogen.

### *In vitro* antiplasmodial activity assay

Compounds were tested at different highest final concentration (between 12.5 µM and 350 µM) depending on their solubility. Stock solutions of the compounds were prepared in dimethyl sulfoxide (DMSO) or water, followed by dilution in RPMI 1640 medium in the absence of thiamine (formulated in-house from the individual components of RPMI 1640, excluding thiamine) or in the presence of 2.97 µM (the concentration normally present in RPMI 1640) or 297 µM thiamine. The final concentration of DMSO that the parasites were exposed to never exceeded 0.05%, a concentration that has no effect on parasite proliferation (23). Four analogues (fursultiamine, cycotiamine, allithiamine, and ethyl thiamine) in addition of oxythiamine and N3PT **9** were further investigated for their antiplasmodial activity against *P. knowlesi* in the presence of 2.97 μM thiamine. Two-fold serial dilutions were performed, with each concentration tested in triplicate. The assay was performed as described, with some modifications (24). Experiments were initiated with parasites in the ring-stage with a parasitemia of 0.5% (*P. falciparum*) and mixed-stage with a parasitemia level of 1% (*P. knowlesi*) and a haematocrit of 1%. For experiments designed to allow comparison of the activity of the compounds against *P. falciparum* and *P. knowlesi*, asynchronous *P. falciparum* cultures were used to mimic those of *P. knowlesi*. Chloroquine (0.5 µM) was used as the positive control (*i.e.* complete inhibition of parasite proliferation), and parasites maintained in the absence of any inhibitor represented 100% parasite proliferation. The final volume in each well was 200 µL. Plates were incubated at 37 °C under an atmosphere of 96% nitrogen, 3% carbon dioxide and 1% oxygen.

Parasite proliferation was measured using the SYBR-Safe assay (25), which correlates fluorescence intensity to parasite DNA. Oxythiamine, fursultiamine, and allithiamine appeared to be incompatible with this fluorescence-based assay (there were inconsistent fluorescence intensity readings at some concentrations). For these compounds, the malstat assay was therefore used instead. The malstat assay correlates parasite lactate dehydrogenase activity with parasite proliferation during the incubation period (96 h for *P. falciparum* and 54 h for *P. knowlesi*) (26). The concentration at which the compound suppresses parasite proliferation by 50% (IC_50_) was determined from non-linear regression plots using GraphPad Prism. The data were averaged from three independent experiments.

### Generation of parasites expressing *Pf*TPK-GFP

The generation of parasites expressing a copy of *Pf*TPK tagged to GFP, in addition to the endogenous *Pf*TPK was described previously (20). The transgenic parasites were maintained under WR99210 (10 nM) pressure.

### Cytotoxicity evaluation of selected compounds

Cytotoxicity testing of selected compounds was conducted using human foreskin fibroblasts (HFF) as described (27), with some modifications. Dulbecco’s Modified Eagle’s Medium (DMEM) with 10% heat-inactivated fetal bovine serum was used for the assay. Briefly, the HFF cells were seeded in 96-well plates at a density of about 25 × 10^4^ cells/mL. Cycloheximide (10 µM; a protein synthesis inhibitor) was used as a control to indicate complete inhibition of HFF cell proliferation. Plates were incubated at 37°C in a humidified, 5% carbon dioxide incubator for 96 h. A sample of the supernatant (150 µL) was then carefully aspirated from each well and discarded. The plates were then stored at -80°C. SYBR-Safe assay was used to measure cell proliferation. Briefly, the plates were thawed, SYBR-Safe lysis solution (150 µL) was added to each well and mixed *via* pipetting to ensure the HFF cells were detached from the plate and lysed. The plates were then processed as described for the antiplasmodial assay.

### Radioactive thiamine uptake assay

[^3^H]Thiamine accumulation (a combination of thiamine transport and its metabolism into TPP) was measured using a method previously applied for other vitamins (28, 29), with some modifications. *P. falciparum* trophozoites were isolated from erythrocytes using 1% saponin. The isolated parasites were washed twice in 50 mL malaria saline (125 mM NaCl, 5 mM KCl, 25 mM HEPES, 20 mM glucose, 1 mM MgCl_2_, pH adjusted to 7.1). On average, the number of cells used was 0.8 × 10^8^ cells/mL. [^3^H]Thiamine, with a final concentration of 0.2 µCi/mL (200 nM), was added to the cell suspension. At predetermined time points (over 30 min), 200 µL of the mixture was added to an oil mix (a 5:4 mix of dibutyl phthalate:dioctyl phthalate). The mixture was centrifuged, washed three times, lysed with 0.1% triton, and precipitated with 5% trichloroacetic acid. A 20 µL aliquot of the supernatant of the first time point was taken to determine the concentration of radioactivity in the extracellular solution. The uptake of [^3^H]thiamine was expressed as CPM_in_/CPM_out_, defining the concentration of [^3^H]thiamine (and [^3^H]TPP generated by the parasite) inside the cell relative to that outside of the cell.

### Computational docking

The docking of N3PT **9**, oxythiamine **3** and thiamine **1** to the *Pf*TPK AlphaFold model (AF-Q14RW4-F1) was performed using the CCDC GOLD docking program. To identify the binding pockets and generate a holoenzyme, the model was superimposed onto a holoenzyme of mouse TPK (PDB: 2f17), with a metal ion and products of TPK (AMP and pyrithiamine pyrophosphate, an analogue of thiamine pyrophosphate that had been used in the crystal structure determination). The thiamine pyrophosphate binding site was subsequently used to dock **1**, **3** and **9** while the AMP binding site was used to dock ATP. ATP, **1**, **3** and **9** were generated using CCDC Mercury. The Genetic Algorithm (GA) runs were conducted with a set of 50 runs. Early termination was not allowed during these runs. To replicate their binding positions, the compounds were subjected to similarity and scaffold constraints based on the original ligands. The docking scoring and rescoring methods employed for analysis were CHEMPLP and GoldScore respectively (30, 31). For molecular docking using CCDC GOLD, the best-performing ligand from each compound series was chosen as a representative.

### *In vivo* antimalarial activity assay

Female Swiss mice (8 weeks old) were used for this experiment. The study was conducted in strict accordance with the German ’Tierschutzgesetz in der Fassung vom 22. Juli 2009’ and Directive 2010/63/EU of the European Parliament and Council, which focuses on the protection of animals used for scientific purposes. The protocol received approval from the ethics committee of the Berlin state authority (‘Landesamt für Gesundheit und Soziales Berlin’, permit number [G029415]). To assess the effectiveness of N3PT **9** in suppressing parasite proliferation in mice, Peter’s 4-day suppression test was used (32). To induce infection, naive mice were injected intravenously with 1 × 10^4^ *P. berghei* parasites, ANKA strain, obtained from an infected donor mouse. The mice were then divided into three groups, each consisting of three individuals. Two hours post-infection, one group received 100 mg/kg and a second group received 200 mg/kg of N3PT **9** dissolved in 100 µL phosphate buffered saline (PBS) *via* intraperitoneal (IP) injection. The third, group of infected mice served as the control and were administered with 100 µL PBS as a vehicle control. Repeat doses were given at intervals of 24, 48, and 72 h after the initial dose. Throughout the duration of the experiment, any signs of toxicity, such as changes in movement and body weight, were monitored and recorded daily. Forty-eight hours after the final drug administration, a small amount of blood was collected from the tail of each mouse to prepare Giemsa-stained slides. The parasitemia of the mice was determined by counting a minimum of 500 erythrocytes. The counting was carried out in a blinded fashion – which parasitemia belonged to which group was only revealed once the counting had been complete.

### Statistical Analysis

Statistical differences were determined using paired or unpaired, two-tailed Student’s t-tests, as appropriate.

## RESULTS

### *In vitro* antiplasmodial activity and cytotoxicity

Twelve thiamine analogues, oxythiamine **3** and **9**-**19,** were tested against the chloroquine-sensitive strain (3D7) of *P. falciparum* in the absence or presence of different extracellular thiamine concentrations (Figure S1). Alongside oxythiamine **3** (consistent with previous observations (15)), N3PT **9**, fursultiamine **10**, allithiamine **11**, cycotiamine **12**, and ethyl thiamine **13** possessed antiplasmodial activity at the concentrations tested (Table 1). N3PT **9** exhibited significantly more potent inhibition of parasite proliferation compared to oxythiamine **3** at all the concentrations of **1**: 14-fold lower IC_50_ in thiamine-free medium (*p* = 0.04, paired t-test), 62-fold lower in the presence of 2.97 µM thiamine (*p* = 0.0006, paired t-test), and 58-fold lower in the presence of 297 µM thiamine (*p* = 0.001, paired t-test). Increasing the extracellular thiamine concentration from 0 to 297 µM reduced the antiplasmodial activity of both **3** and **9,** but to different degrees, with oxythiamine **3** showing a 986-fold reduction in activity and N3PT **9** having its activity reduced 220-fold (Table 1 and Figure S2). However, increasing the extracellular thiamine concentration has no effect on the antiplasmodial activity of compounds **10-13**, consistent with them acting *via* a mechanism unrelated to thiamine utilization. Compounds **9-13** were then tested against *P. knowlesi*, a surrogate of *P. vivax* (33), the second most important *Plasmodium* species in terms of morbidity and mortality (34). All the compounds except for **13** inhibited the proliferation of *P. knowlesi*, albeit with IC_50_ values, in general, being higher than those against *P. falciparum* (Table 1 and Figure S2). They were then also tested against HFF cells to determine if they are cytotoxic. While **10** and **11** showed low micromolar activity against HFF cells, compounds **9**, **12** and **13** were not cytotoxic (Figure S3).

**Table 1.**
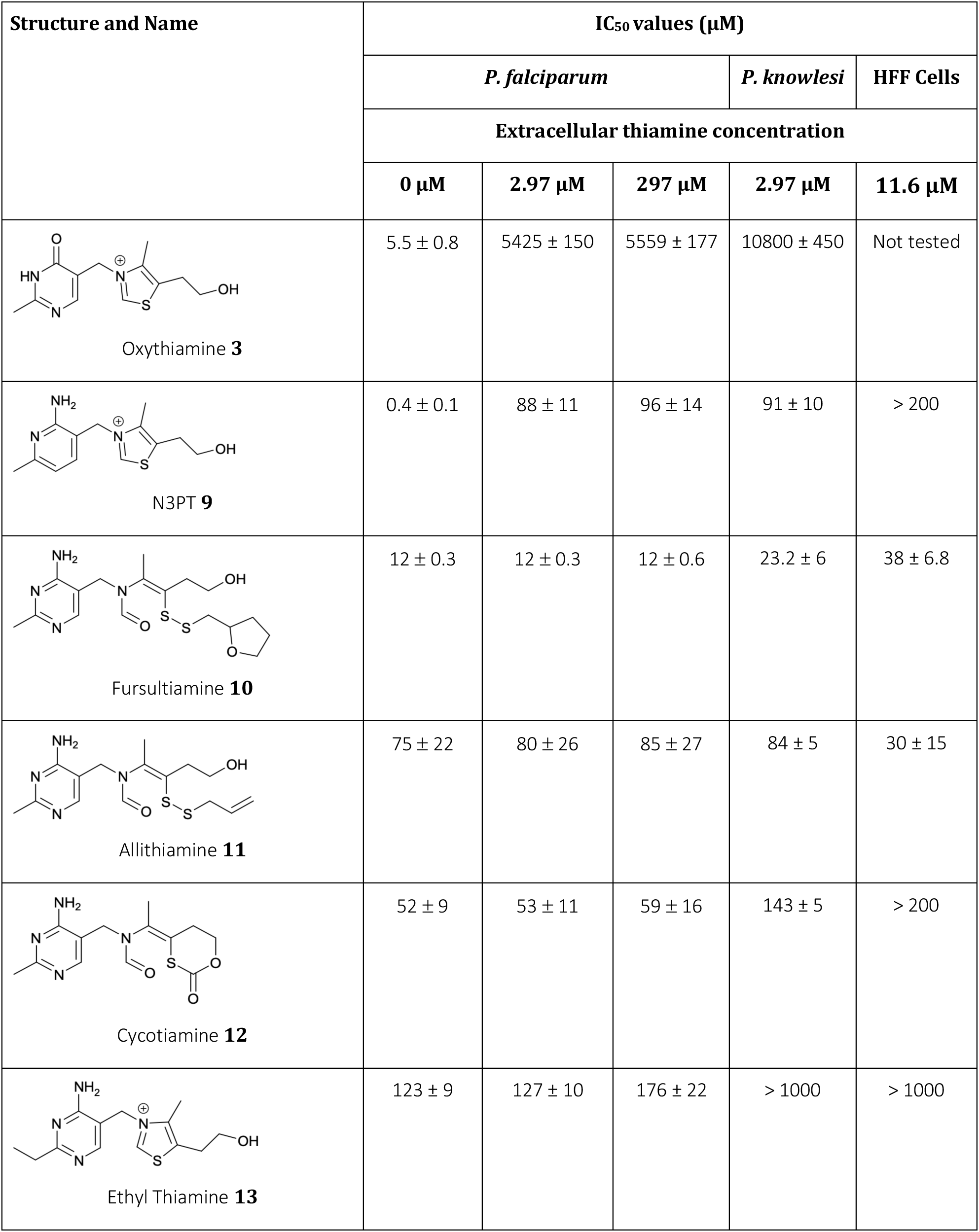
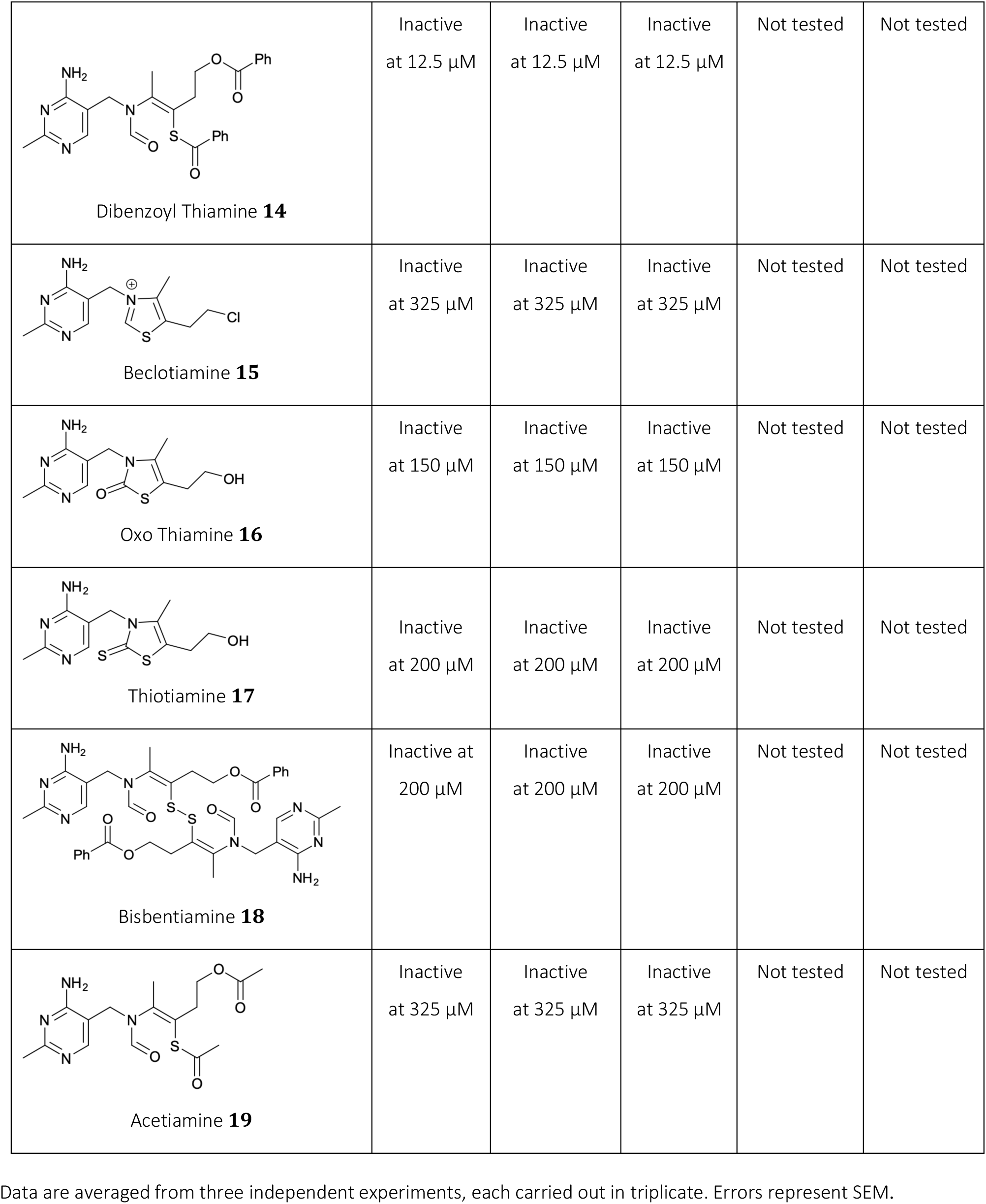
Antiplasmodial activity of the thiamine analogues against *P. falciparum* and *P. knowlesi*, and their cytotoxicity against HFF cells.

| Structure and Name | IC<sub>50</sub> values (μM) |  |  |  |  |
| --- | --- | --- | --- | --- | --- |
|  | *P. falciparum* |  |  | *P. knowlesi* | HFF Cells |
|  | Extracellular thiamine concentration |  |  |  |  |
|  | 0 μM | 2.97 μM | 297 μM | 2.97 μM | 11.6 μM |
| Oxythiamine **3** | 5.5 ± 0.8 | 5425 ± 150 | 5559 ± 177 | 10800 ± 450 | Not tested |
| N3PT **9** | 0.4 ± 0.1 | 88 ± 11 | 96 ± 14 | 91 ± 10 | > 200 |
| Fursultiamine **10** | 12 ± 0.3 | 12 ± 0.3 | 12 ± 0.6 | 23.2 ± 6 | 38 ± 6.8 |
| Allithiamine **11** | 75 ± 22 | 80 ± 26 | 85 ± 27 | 84 ± 5 | 30 ± 15 |
| Cycotiamine **12** | 52 ± 9 | 53 ± 11 | 59 ± 16 | 143 ± 5 | > 200 |
| Ethyl Thiamine **13** | 123 ± 9 | 127 ± 10 | 176 ± 22 | > 1000 | > 1000 |
| Dibenzoyl Thiamine **14** | Inactive<br>at 12.5 μM | Inactive<br>at 12.5 μM | Inactive<br>at 12.5 μM | Not tested | Not tested |
| Beclotiamine **15** | Inactive<br>at 325 μM | Inactive<br>at 325 μM | Inactive<br>at 325 μM | Not tested | Not tested |
| Oxo Thiamine **16** | Inactive<br>at 150 μM | Inactive<br>at 150 μM | Inactive<br>at 150 μM | Not tested | Not tested |
| Thiotiamine **17** | Inactive<br>at 200 μM | Inactive<br>at 200 μM | Inactive<br>at 200 μM | Not tested | Not tested |
| Bisbentiamine **18** | Inactive at<br>200 μM | Inactive<br>at 200 μM | Inactive<br>at 200 μM | Not tested | Not tested |
| Acetiamine **19** | Inactive<br>at 325 μM | Inactive<br>at 325 μM | Inactive<br>at 325 μM | Not tested | Not tested |

Data are averaged from three independent experiments, each carried out in triplicate. Errors represent SEM.

### Effect of TPK overexpression on parasite sensitivity to thiamine analogues

To determine whether the mode of antiplasmodial action is dependent on TPK, compounds were tested against a *P. falciparum* cell line (termed *Pf*TPK-GFP), which expresses a GFP-tagged copy of TPK, in addition to the endogenous TPK. In both thiamine-free and thiamine-replete conditions, the *Pf*TPK-GFP cell line was hypersensitive to oxythiamine **3** when compared to control parasites transfected with a plasmid expressing only GFP (termed *Pf*3D7-GFP; Figure 2), consistent with previous observations (15). Having confirmed that the *Pf*TPK-GFP cell line was behaving as expected, a few analogues (**9, 10 and 12)** were selected, based on their antiplasmodial activity and structural modifications, and tested against these parasites. Fursultiamine **10** and cycotiamine **12** were equally active against the *Pf*TPK-GFP cell line as they were against the control parasites (Figure S4), consistent with TPK not playing a role in the antiplasmodial activity of these compounds. In contrast, and as observed previously for oxythiamine **3** (15), N3PT **9** was more active against the *Pf*TPK-GFP cell line under thiamine-free conditions (IC_50_ = 0.05 ± 0.02 µM) when compared to *Pf*3D7-GFP parasites (IC_50_ = 0.36 ± 0.03 µM, *p* = 0.002, unpaired t-test), although the effect was somewhat less pronounced than that observed with oxythiamine **3** (7-fold versus 13-fold) (Figure 2). However, in the presence of 297 µM thiamine, the potency of N3PT **9** against both parasite lines was comparable (*Pf*TPK-GFP: IC_50_ = 93 ± 4 µM; *Pf*3D7-GFP: IC_50_ = 98 ± 2 µM, *p* = 0.13, unpaired t-test), unlike what we observed with oxythiamine **3** (*Pf*TPK-GFP: IC_50_ = 275 ± 59 µM; *Pf*3D7-GFP: IC_50_ = 8220 ± 267 µM, *p* = 0.0004, unpaired t-test) (Figure 2).

**Figure 2.**
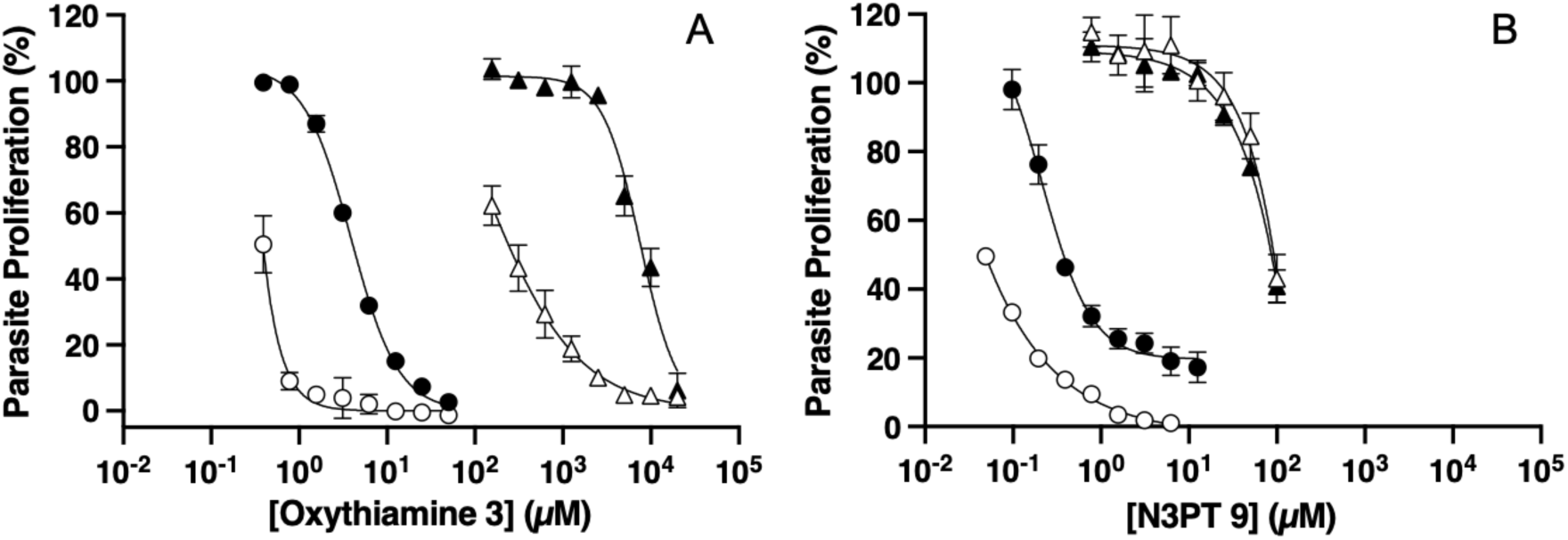
Antiplasmodial activity of oxythiamine **3** (A) and N3PT **9** (B) against *Pf*TPK-GFP (white symbols) and *Pf*3D7-GFP (black symbols). The experiments were carried out in thiamine-free medium (circles) or medium containing 297 µM thiamine (triangles). Data are averaged from three independent experiments, each carried out in triplicate. Error bars represent SEM and, where not visible, are smaller than the symbols.

### The effect of 3 and 9 on [^3^H]thiamine accumulation

A [^3^H]thiamine accumulation assay was conducted to investigate the effects of compounds **3** and **9** on thiamine uptake into the parasite and its subsequent metabolism into TPP **2** (and any other thiamine-derived metabolites). In the absence of the analogues, isolated parasites accumulated [^3^H]thiamine and thiamine metabolites at a rate of 0.8 ± 0.3 CPM_in_/CPM_out_ *per* min (measured in the linear part of the time course, 5-30 min). The presence of 100 µM N3PT **9** and a higher concentration of oxythiamine **3** (500 µM; chosen due to its lower antiplasmodial potency when compared to **9**), significantly reduced [^3^H]thiamine/metabolite accumulation in isolated parasites by more than 3-fold (0.2 ± 0.04, and 0.21 ± 0.02 CPM_in_/CPM_out_ *per* min, respectively, p ::; 0.04, paired t-test, Figure 3). These data are consistent with compounds **3** and **9** reducing [^3^H]thiamine/metabolite accumulation by competing with **1** for thiamine transporters and/or metabolism by TPK.

**Figure 3.**
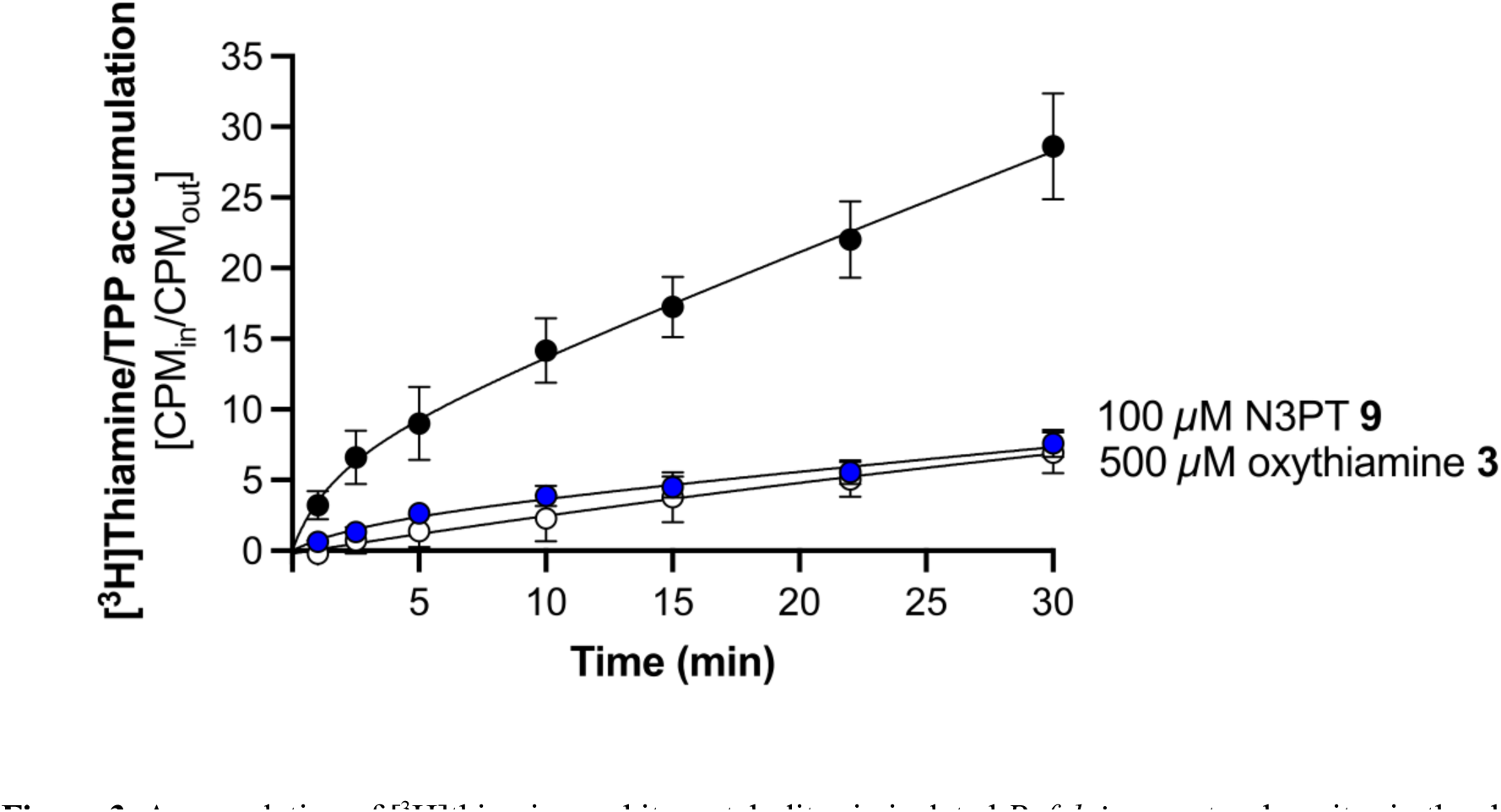
Accumulation of [^3^H]thiamine and its metabolites in isolated *P. falciparum* trophozoites in the absence (black circles) or presence of 100 µM N3PT **9** (blue circles), or 500 µM oxythiamine **3** (white circles). Data are averaged from three independent experiments, each carried out in duplicate. Error bars represent SEM and where not visible, are smaller than the symbols.

### Docking of oxythiamine 3 and N3PT 9 into the active site of *Pf*TPK

Since N3PT **9** only has a small structural modification in the aminopyrimidine ring when compared to thiamine, computational docking was performed to determine whether it is likely to bind in the active site of *Pf*TPK, supporting the data observed in Figure 3. With no structure of *Pf*TPK available from either crystallography or cryo-EM, an AlphaFold model (AF-Q14RW4-F1) was used. The model is an apoenzyme consisting of only the tertiary structure of the protein. To identify the binding pockets and generate a holoenzyme with its substrate, the model was superimposed onto a holoenzyme of mouse TPK (PDB: 2f17), which has a sequence identity with *Pf*TPK of 33.5%, and has a metal ion, AMP and pyrithiamine pyrophosphate, an analogue of thiamine pyrophosphate, all bound (35) (Figure 4A). The thiamine pyrophosphate binding site was subsequently used to dock thiamine **1,** oxythiamine **3** and N3PT **9** while AMP was docked into the AMP pocket (Figures 4B-D). N3PT **9** has been shown to be converted into N3PT pyrophosphate by the human TPK resulting in inhibition of transketolase activity in human colon cancer cells (36). Similarly, oxythiamine 3 is a known substrate of *Pf*TPK (15), and it is oxythiamine pyrophosphate that is likely toxic to parasites. The docking results (Figure 4) showing that N3PT 9, thiamine and oxythiamine 3 have very similar binding poses (Figure 4B-D and Figure S5), together with the latter two compounds being known substrates of *Pf*TPK, supports the hypothesis that N3PT **9**is also a substrate of *Pf*TPK. The docked model of thiamine has one hydrogen bond from N-1 to the OH of Ser371 and one from the 4-NH_2_ group to the main chain C=O of Gln257. However, it seems likely that the side-chain of Glu258 would twist round to form a second hydrogen bond to the NH_2_ group – this is what happens in PDB: 2f17, which has an Asp at the equivalent position. Compared to thiamine’s pyrimidine ring, N3PT 9 has a pyridine ring (N-1 replaced by CH), whereas oxythiamine 3 has an oxopyrimidine in place of thiamine’s aminopyrimidine (NH_2_ replaced by O). These structural differences would mean that N3PT 9 would have one less hydrogen bond interaction with *Pf*TPK when compared to thiamine, and oxythiamine 3 would have one or two less hydrogen bonds. This would mean that 9 and 3 are likely to have lower affinities than thiamine, and 3 may have lower affinity than 9.

**Figure 4.**
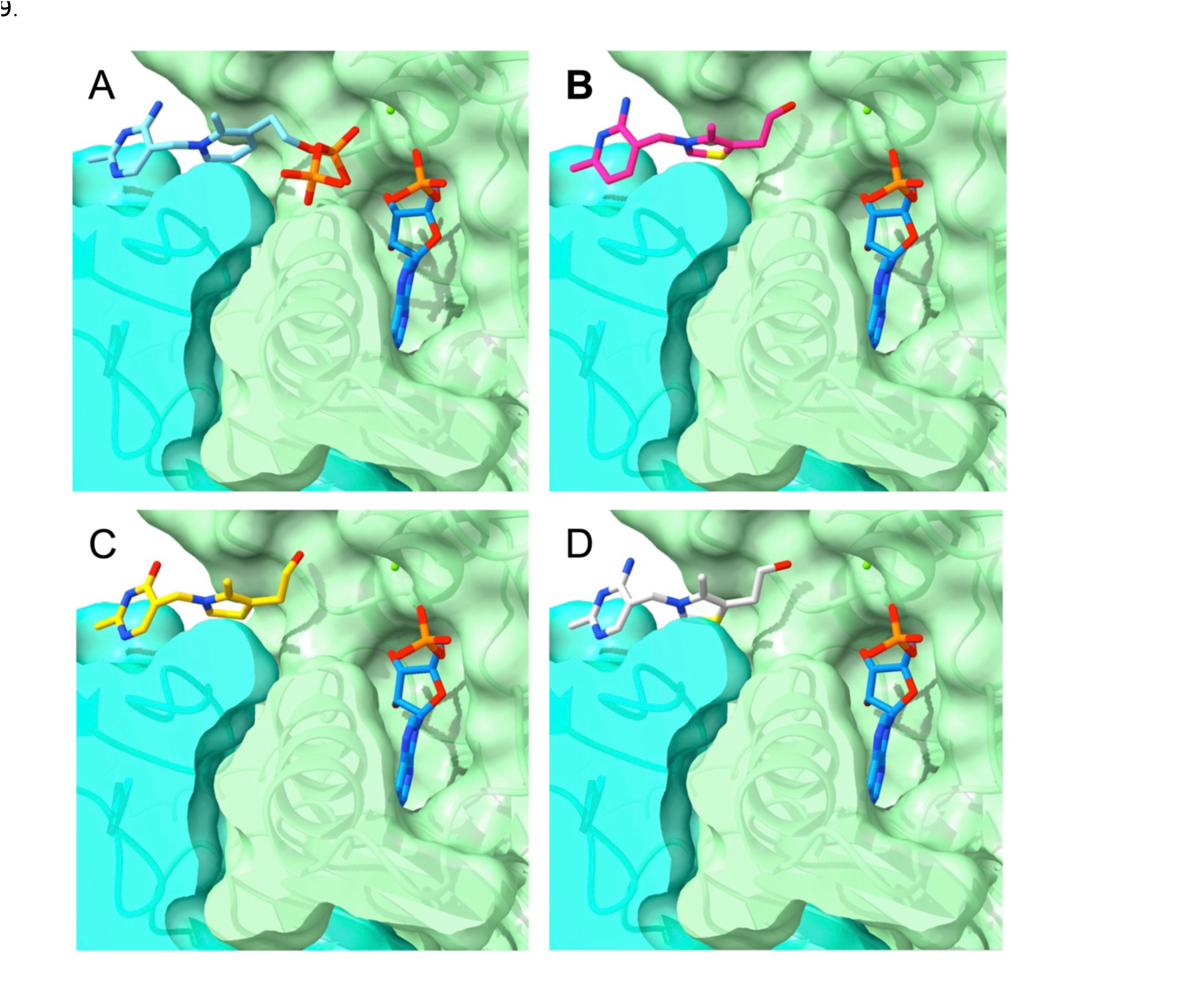
Docking of various compounds into the active site of *Pf*TPK (shown as surface and ribbon) based on an AlphaFold structure (AF-Q14RW4-F1) modelled on the crystal structure of the mouse TPK (PDB: 2f17). Cyan and green colours represent the two protomers of the putative *Pf*TPK dimer. *Pf*TPK with (A) the ligands in the mouse TPK crystal structure, pyrithiamine pyrophosphate and AMP, (B) dockings of AMP and N3PT **9** in pink, (C) oxythiamine **3** in yellow, and (D) thiamine **1** in white.

### *In vivo* antimalarial activity of N3PT 9

To assess whether N3PT **9** possesses antimalarial activity *in vivo*, we tested it in the murine malaria model, using the *P. berghei* ANKA strain and Peter’s Suppressive Test protocol. Two doses of **9** were tested, 100 and 200 mg/kg, administered intraperitoneally. Signs of toxicity, such as weight loss and reduced movement were monitored from day 0 to day 5 post-infection (Figure S6). N3PT **9** showed a dose-dependent reduction in the parasitemia of infected mice (*p* = 0.0015 and *p* < 0 .0001 for the 100 and 200 mg/kg doses, respectively; Student’s t-test) relative to the non-treated group (vehicle only; Figure 5A). Although the 100 mg/kg dose did not increase the time to symptoms in mice, the 200 mg/kg dose prolonged their time to symptoms by 9 days compared to the untreated mice (Figure 5B).

**Figure 5.**
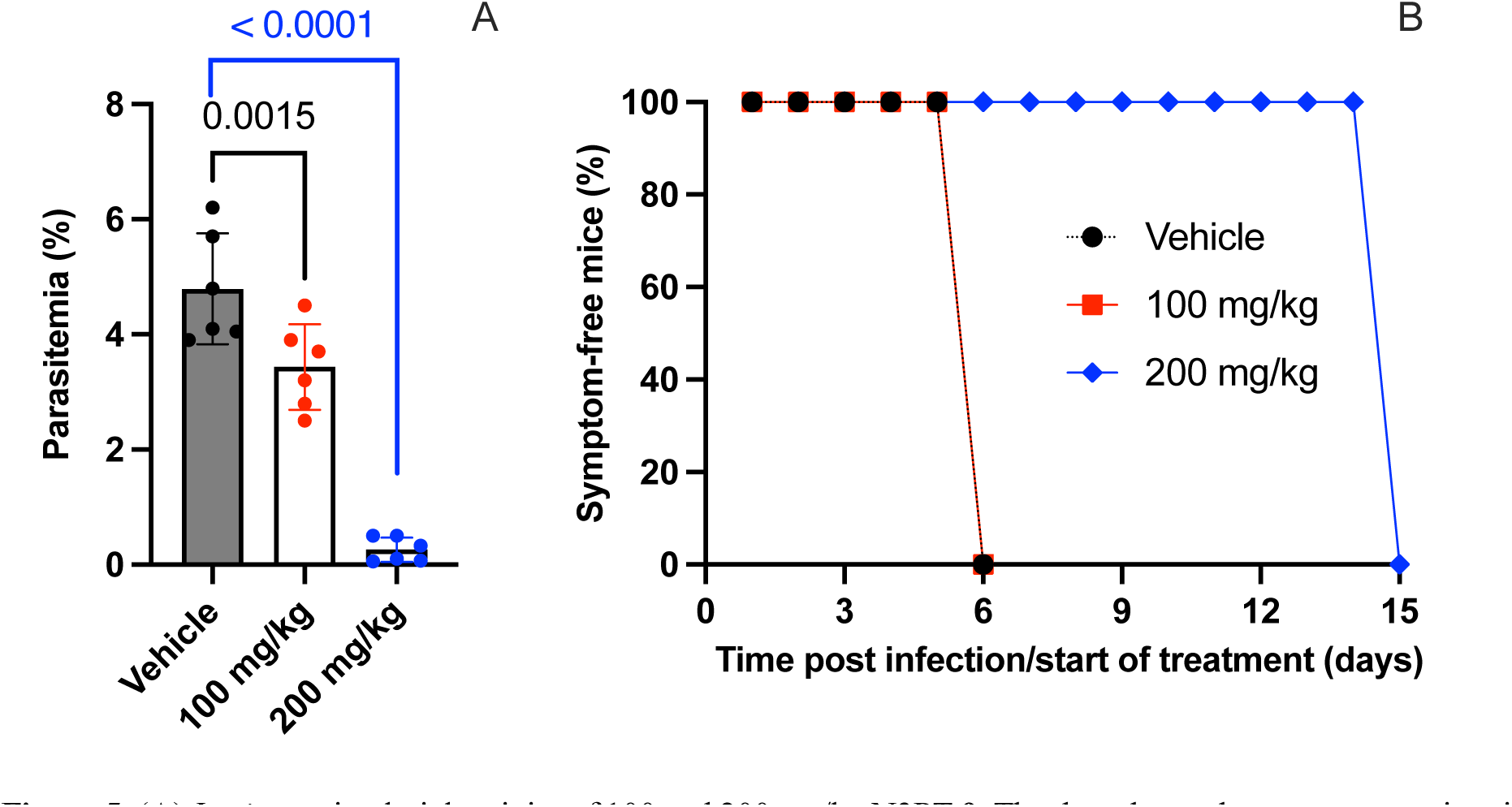
(**A**) *In vivo* antimalarial activity of 100 and 200 mg/kg N3PT **9**. The data shows the average parasitemia at day 5 after infection and daily treatment. Symbols represent data points from individual mice (*n* = 6). Two independent experiments were carried out, each with 3 mice in each group. Bars represent averaged data ±SD. Student’s t-test results are indicated above the bars for relevant comparisons. (**B**) The percentage of symptom-free mice according to treatment: Vehicle/ PBS (black circles), 100 mg/kg N3PT **9** (red squares), and 200 mg/kg (blue diamonds). *n* = 6, from 2 biological replicates.

## Discussion

To identify thiamine analogues with better antiplasmodial activity than oxythiamine **3**, eleven compounds **9-19** were evaluated against two *Plasmodium* species alongside oxythiamine **3**. These twelve compounds together sampled a wide chemical space, with 4/12 being positively charged, 3/12 bearing a modified aminopyrimidine ring, 2/12 bearing a neutral central ring (as for **5-8** that we have previously reported (18–20)), and 6/12 featuring an open scaffold in place of the central ring (Table 1). Compounds **9-13** suppressed *P. falciparum* with IC_50_ values in the micromolar range, whilst **14-19** were unable to inhibit parasite proliferation by more than 50% at the highest concentration tested (which varied depending on the solubility of the compound). Compounds **13** and **15**, which differ from thiamine’s pyrimidine ring and the hydroxyl tail respectively, may be able to hijack the thiamine transporter for cell entry due to their overall structural similarity. While both compounds may compete with intracellular thiamine for binding to TPK, only ethyl thiamine **13** can be activated into **13**-pyrophosphate (**13**-PP), which in turn competes with TPP for the TPP-dependent enzymes’ coenzyme pocket. Oxo thiamine **16** and thiotiamine **17** featuring a neutral central ring (as with **7**) failed to inhibit parasite proliferation by more than 30% at the highest concentration tested, probably because they do not have a proper metal-binding group on their tail to enable efficient enzyme binding (18–20). Compounds **10**-**12**, **14**, **18** and **19** are all precursors of thiamine and because they are neutral, they may be able to enter cells *via* simple diffusion. In the cytoplasm, they are chemically transformed into the open-thiol species **20** (Figure 6), which in turn spontaneously cyclizes into thiamine. They can, therefore, function as thiamine supplements. Because fursultiamine **10** and allithiamine **11** have greater oral bioavailability than the thiamine salt, they have been used to treat beriberi (a disease caused by thiamine deficiency) (37). Upon reduction, fursultiamine **10** and allithiamine **11** produce a corresponding thiol by-product alongside **20**. The observed antiplasmodial effect of these compounds can neither be antagonized by increasing the extracellular thiamine concentration nor can the antiplasmodial activity of fursultiamine **10** be altered by expression of *Pf*TPK-GFP. Accordingly, the inhibitory effect may be due to a non-specific action of their released thiol, which in turn may lead to cell damage, as intracellular thiol oxidation produces hydrogen peroxide (H_2_O_2_), an oxidizing agent (38). The observed toxic effects in human cells associated with **10** and **11** was a surprising observation given that they have been used in humans as thiamine supplements and may also be attributed to the released thiol. The transformation of dibenzoyl thiamine **14**, bisbentiamine **18** and acetiamine **19** into thiamine generates by-products that are likely to be non-toxic to the parasite at the concentrations used (**14** and **18** produce benzoate and **19** produces acetate). This is consistent with the lack of antiplasmodial activity observed for these compounds. The one exception is cycotiamine **12,** which produces carbonate as a by-product. Carbonate is also expected to be non-toxic to the parasite, so the antiplasmodial action of cycotiamine **12** may be attributed to an off-target mechanism of action by its un-transformed form. This notion is corroborated by our findings that its activity cannot be antagonized by increasing the extracellular concentration of thiamine, nor is it modified by overexpressing *Pf*TPK.

**Figure 6.**
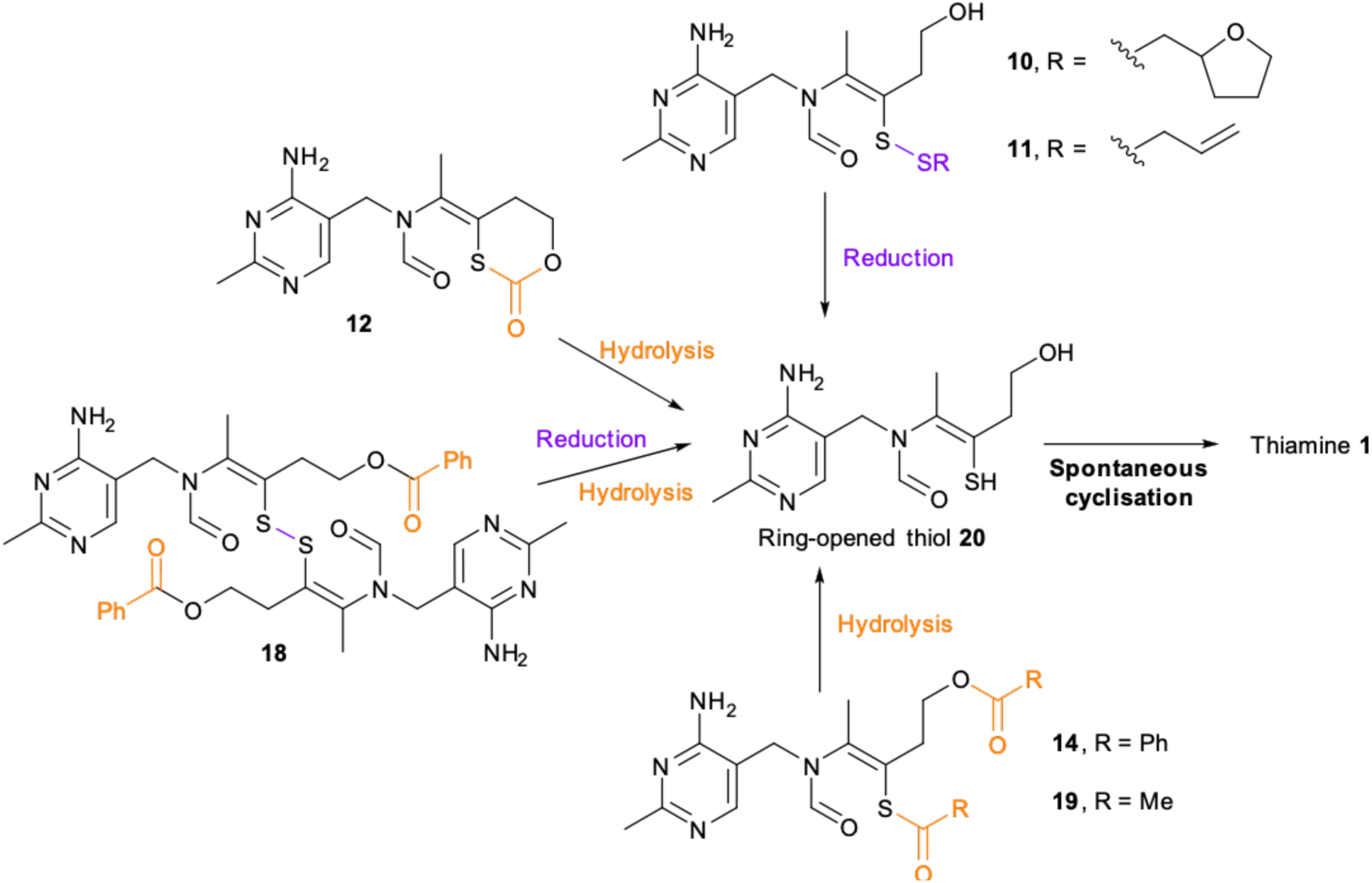
Stepwise transformation of thiamine precursors **10**-**12**, **14**, **18** and **19** through an open-thiol species **20** to thiamine **1**. Substructures of the thiamine precursors that will be released in the course of transformation are coloured. Reduction and hydrolysis are the two main reaction types which are highlighted in purple and orange, respectively.

Upon pyrophosphorylation, N3PT **9** is a known transketolase inhibitor and has shown promising anticancer potential (36, 39). It is the only thiamine analogue we tested that showed sub-micromolar antiplasmodial activity *in vitro* in the absence of extracellular thiamine and was between 10-60 times more potent than oxythiamine **3** (depending on the extracellular thiamine concentration). We observed a >200-fold reduction in antiplasmodial activity of N3PT **9** when thiamine was added to thiamine-free medium (Table 1), demonstrating its competitive relationship with thiamine/TPP. The fact that N3PT **9** and ethyl thiamine **13** are both very similar to thiamine **1** but have very different antiplasmodial activity is consistent with the thiamine utilisation pathway being sensitive to subtle structural changes in the thiamine analogues. Accordingly, ethyl thiamine **13** was not investigated further. N3PT **9** was also active against *P. knowlesi*, had a high selectivity index (Table 1), and was more active than oxythiamine **3**, at half the dose, when tested *in vivo* (Figure 5). This is an important stepping stone towards developing an antimalarial compound targeting this pathway. Synthesis of a range of derivatives of the N3PT **9** lead and systematic *in vitro* testing are warranted to further explore the potential of harnessing the strict dependence of intra-erythrocytic *Plasmodium* replication on thiamine.

While the dependence of oxythiamine **3**’s activation by *Pf*TPK has been established here and previously (15), whether N3PT **9** is also metabolised by *Pf*TPK remains to be shown. To address this, we conducted computational modelling to dock oxythiamine **3**, N3PT **9** and thiamine **1** into the active site of *Pf*TPK.

As shown in Figure 4, the binding mode of N3PT **9** is very similar to that of thiamine **1** and oxythiamine **3**. This is consistent with N3PT **9** being a *Pf*TPK substrate, but the removal of one hydrogen bond from the pyrimidine ring contrasts with the marked higher inhibition of TPK, indicating that for a better understanding of the binding affinity of N3PT **9** to *Pf*TPK structural data are needed. An interaction between N3PT **9** and *Pf*TPK is further supported by the [^3^H]thiamine accumulation experiment, in which [^3^H]thiamine/metabolite accumulation in isolated parasites was significantly reduced in the presence of 100 µM N3PT **9** (Figure 3), consistent with N3PT **9** interacting with TPK, either as a substrate or inhibitor. The same effect was observed with oxythiamine **3,** but at a higher concentration, consistent with its lower antiplasmodial activity. In addition, although in the presence of thiamine the potency of N3PT **9** against parasites expressing *Pf*TPK-GFP is the same as that against control parasites, in thiamine-free medium *Pf*TPK-GFP-expressing parasites are four-fold more sensitive to N3PT **9** when compared to parasites expressing GFP (Figure 2), consistent with its antiplasmodial activity, involving, at least in part, *Pf*TPK.

In conclusion, our data identify N3PT **9** as an improved inhibitor over previously-reported antiplasmodial thiamine analogues. Further medicinal chemistry efforts to optimize its pharmacological properties may afford a novel antimalarial agent targeting parasite thiamine utilization.

## Acknowledgements

We would like to express our gratitude to the van Dooren lab for supplying the HHF cells used in the cytotoxicity assays and Manuel Rauch for his assistance with the *in vivo* experiments. We would also like to thank the Canberra Branch of the Australian Red Cross Lifeblood for the provision of red blood cells. We are grateful to the Developmental Therapeutics Program (DTP) for providing some of the thiamine analogues. IF received support from both the Australian Government Research Training Program (AGRTP) and the German Academic Research Exchange (DAAD). AHYC and TCSH were supported by K.M. Medhealth.

## Author Contributions

IF performed all the biological studies under the supervision of KJS and KM. TCSH and AHYC contributed to the chemistry aspect of the study under the supervision of FJL. IF wrote the first draft of the manuscript with input from KJS, AHYC and TCSH. All authors revised and approved the final version of the manuscript.

## Supplementary Figures

**Figure S1:**
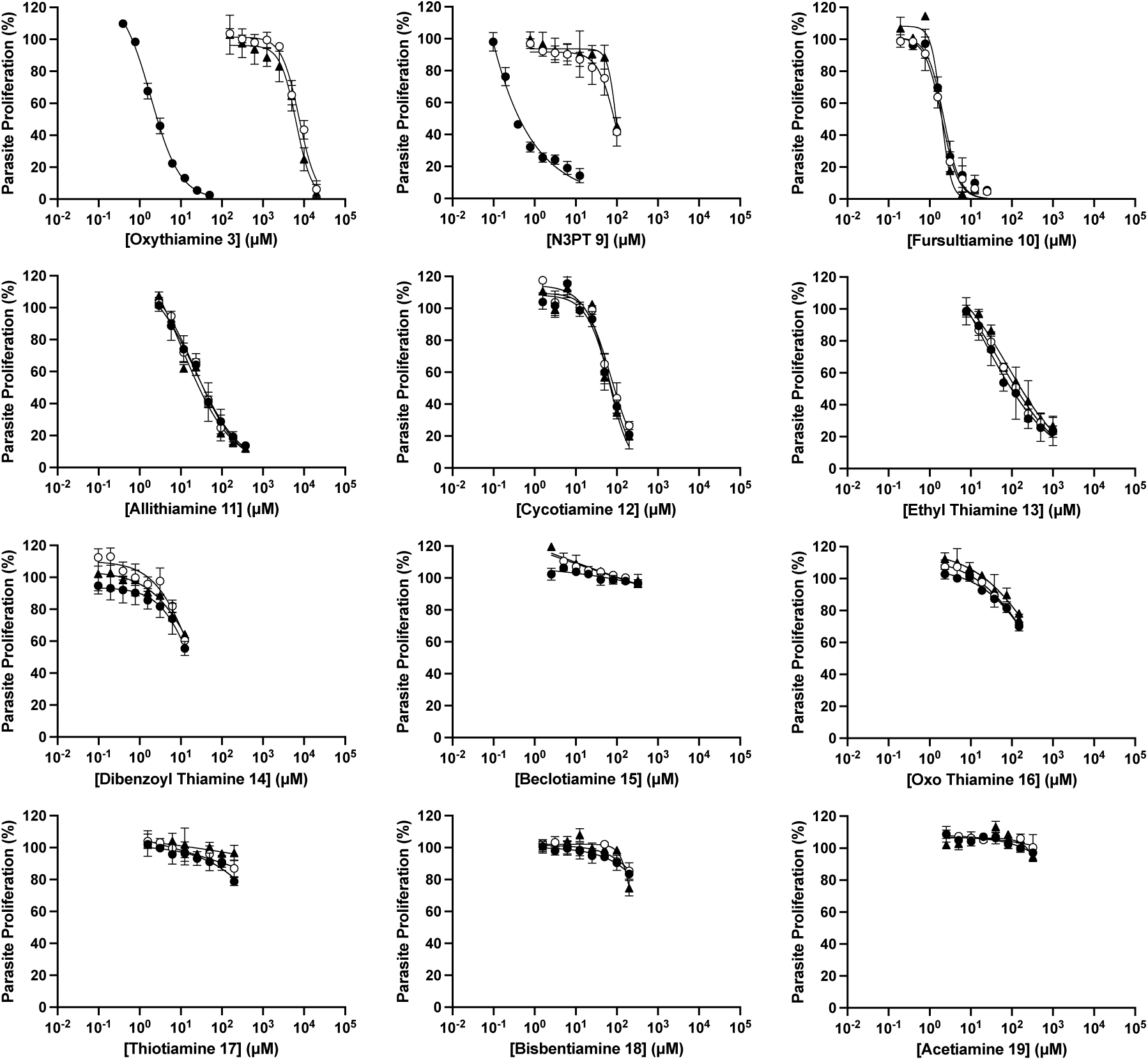
Antiplasmodial activity of thiamine analogues against the *P. falciparum* (strain 3D7) in thiamine-free medium (black circles), medium containing 2.97 µM thiamine (white circles), and medium containing 297 µM thiamine (black triangles). The data are averaged from three independent experiments, each performed in triplicate. Error bars represent SEM, and where not shown, are smaller than the symbol.

**Figure S2:**
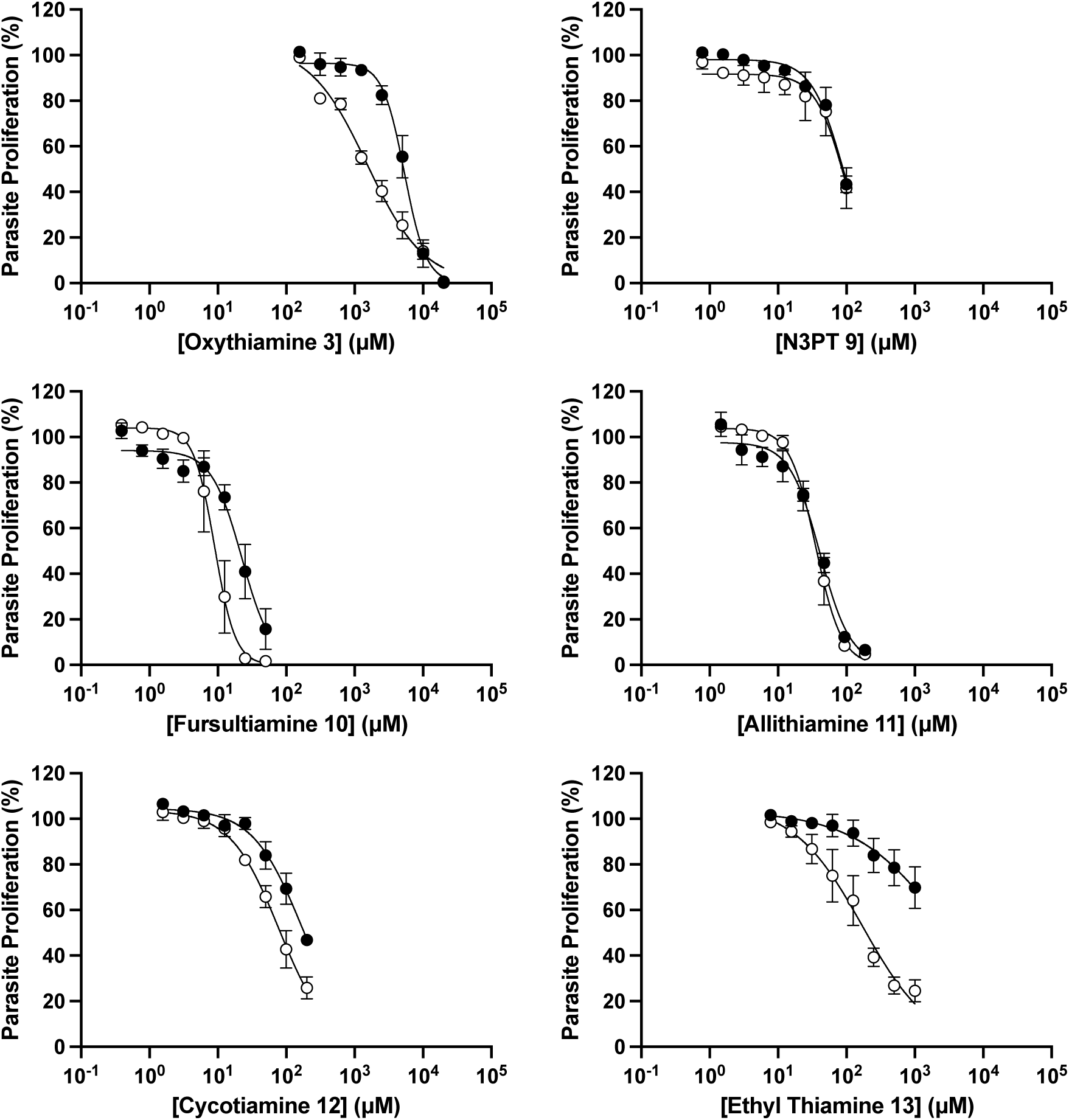
Antiplasmodial activity of oxythiamine 3, N3PT 9, fursultiamine 10, allithiamine 11, cycotiamine 12, and ethyl thiamine 13 against the *P. falciparum* (strain 3D7; white circles), and *P. knowlesi* (strain H1; black circles) in medium containing 2.97 µM thiamine. Note that the experiments with *P. falciparum* were initiated with asynchronous parasites (as opposed to parasites in the ring stage as carried out to generate the data in Figure S1) to match more closely the conditions of the *P. knowlesi* experiments. The data are averaged from three independent experiments, each performed in triplicate. Error bars represent SEM, and where not shown, are smaller than the symbol.

**Figure S3:**
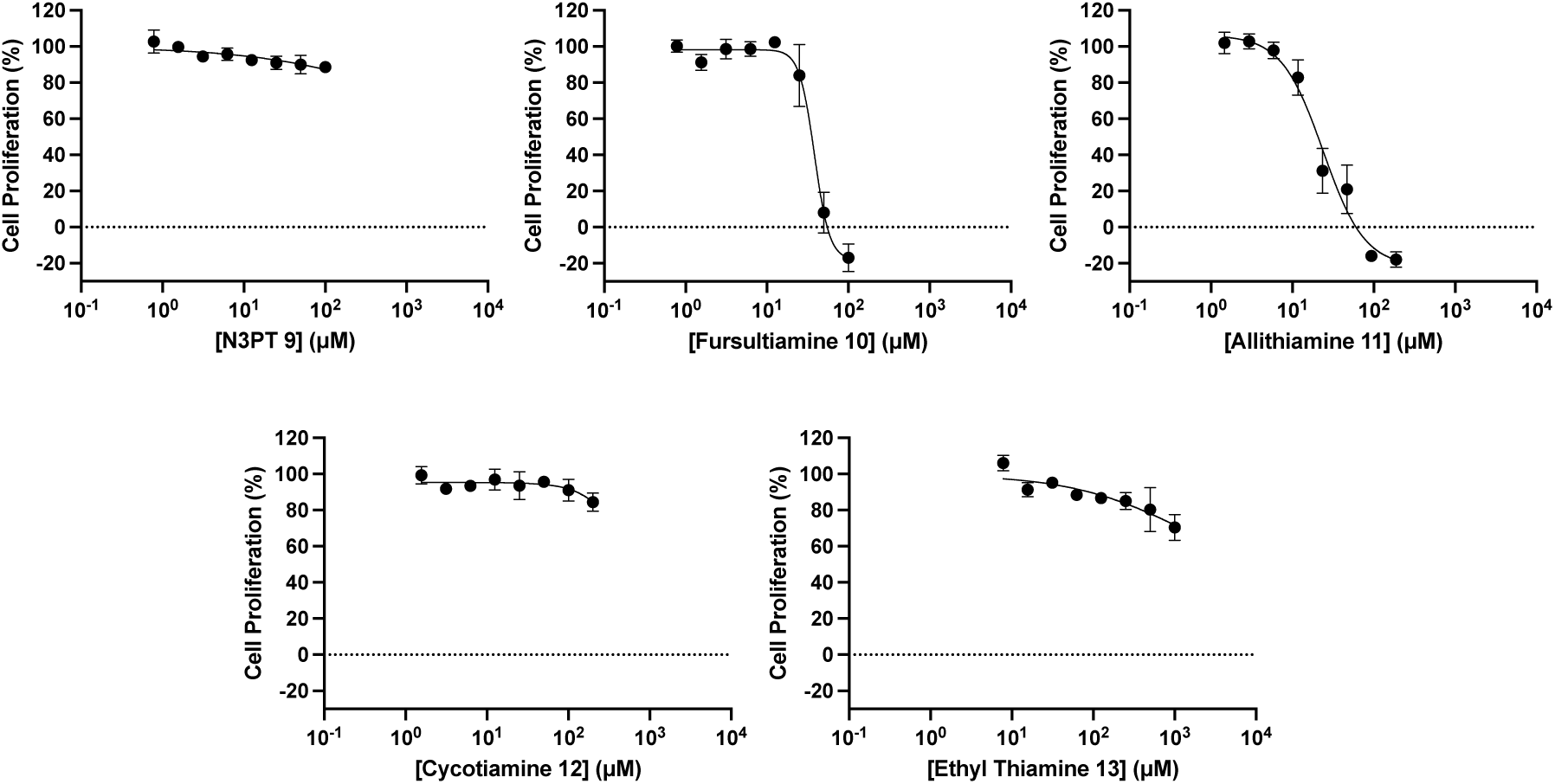
Cytotoxicity of N3PT 9, fursultiamine 10, allithiamine 11, cycotiamine 12, and ethyl thiamine 13 against HFF cells. The data are averaged from three independent experiments, each performed in triplicate. Error bars represent SEM, and where not shown, are smaller than the symbol.

**Figure S4:**
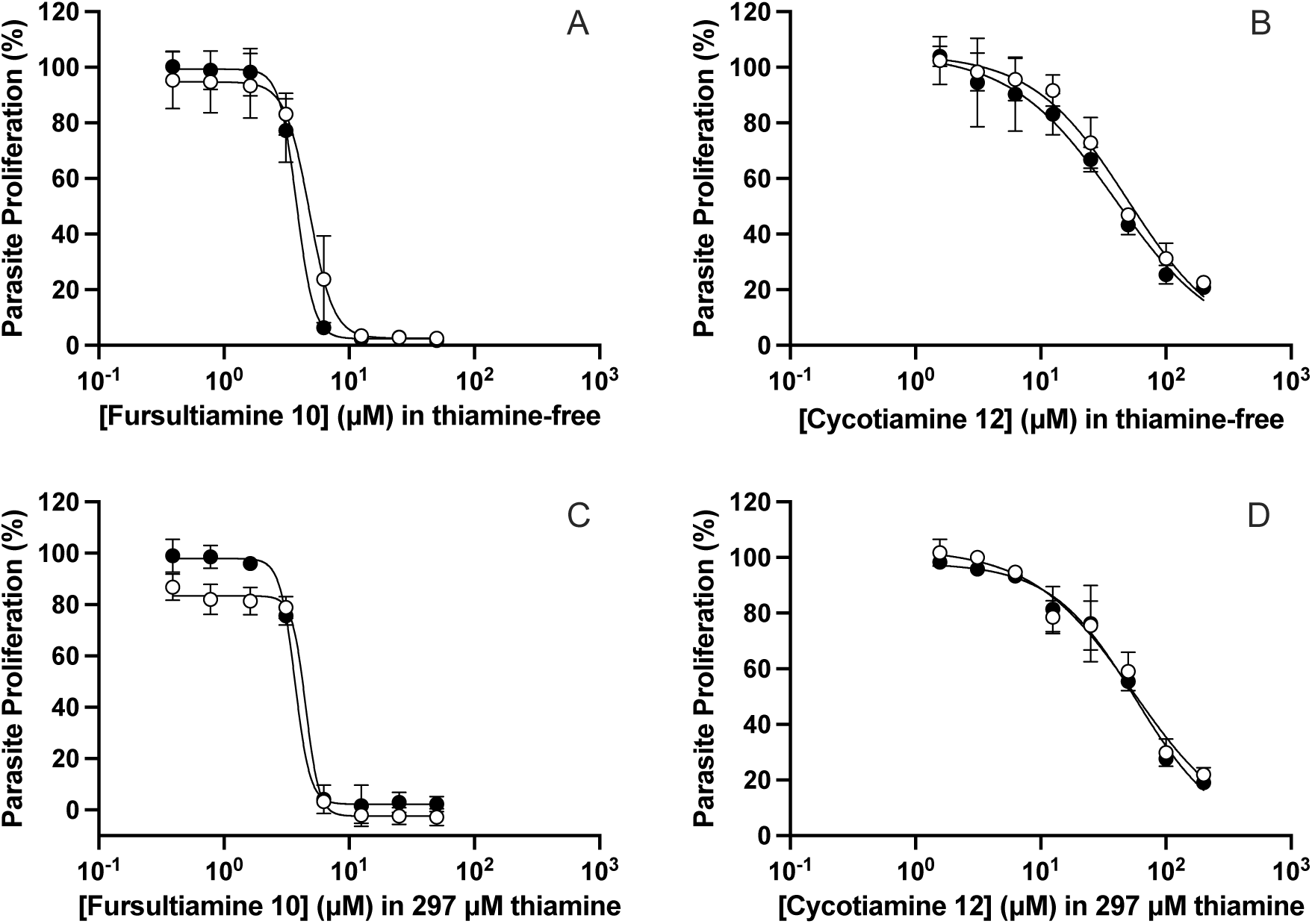
Antiplasmodial activity of fursultiamine 11 and cycotiamine 12 against the *Pf*3D7-GFP (parasites expressing GFP; white circles) and *Pf*TPK-GFP (parasites expressing GFP-tagged TPK, in addition to the endogenous, untagged, TPK; black circles) in thiamine-free medium (A and B) and medium containing 297 µM thiamine (C and D). The data are averaged from three independent experiments, each performed in triplicate. Error bars represent SEM, and where not shown, are smaller than the symbol.

**Figure S5.**
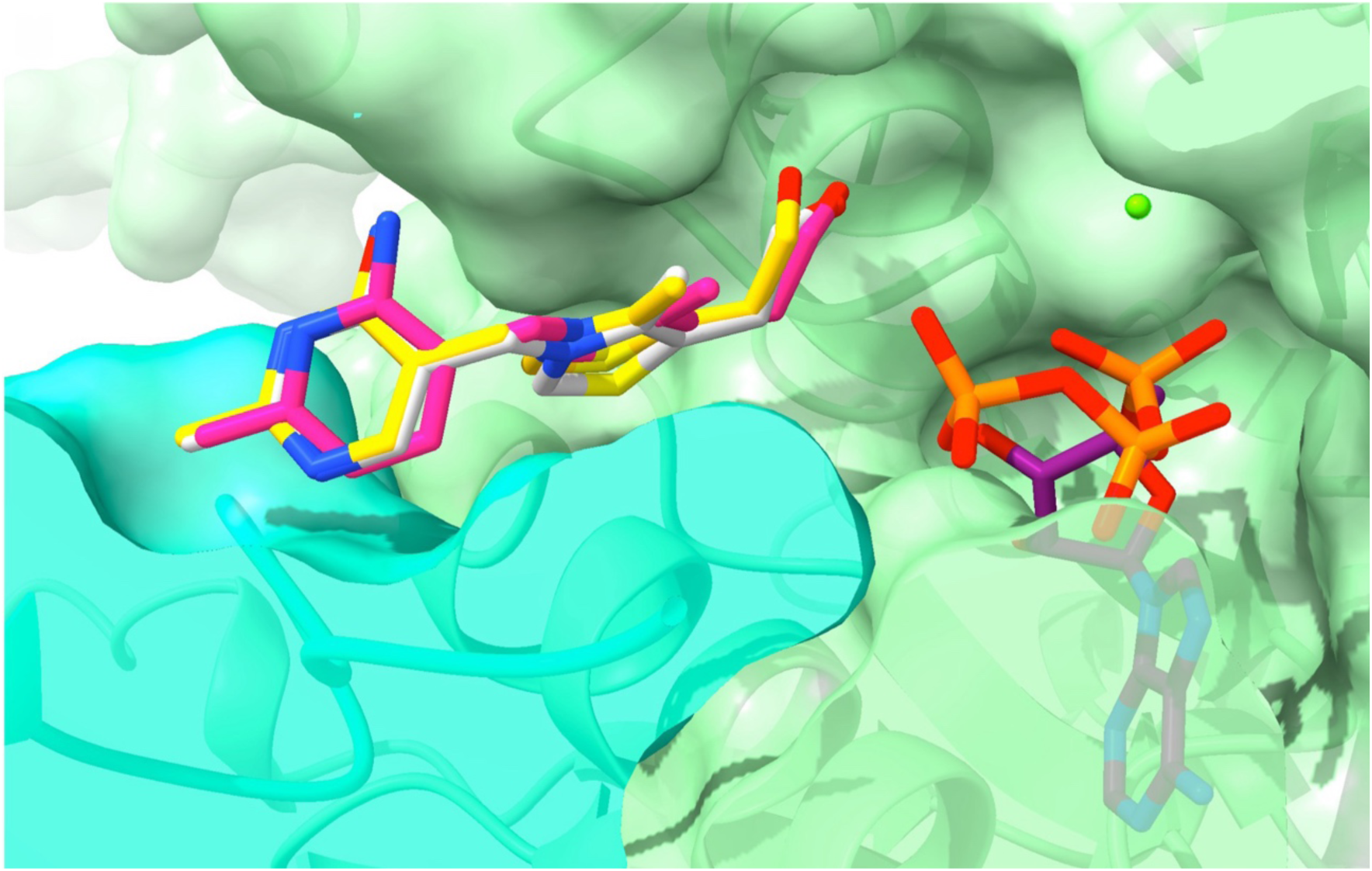
Docking of thiamine and analogues into the active site of *Pf*TPK (shown as surface and ribbon) based on the crystal structure of mouse TPK (PDB: 2f17). ATP is docked into *Pf*TPK, together with N3PT **9** in pink, oxythiamine **3** in yellow and thiamine **1** in white, superimposed on one another in the active site. Magnesium is shown as a green sphere.

**Figure S6:**
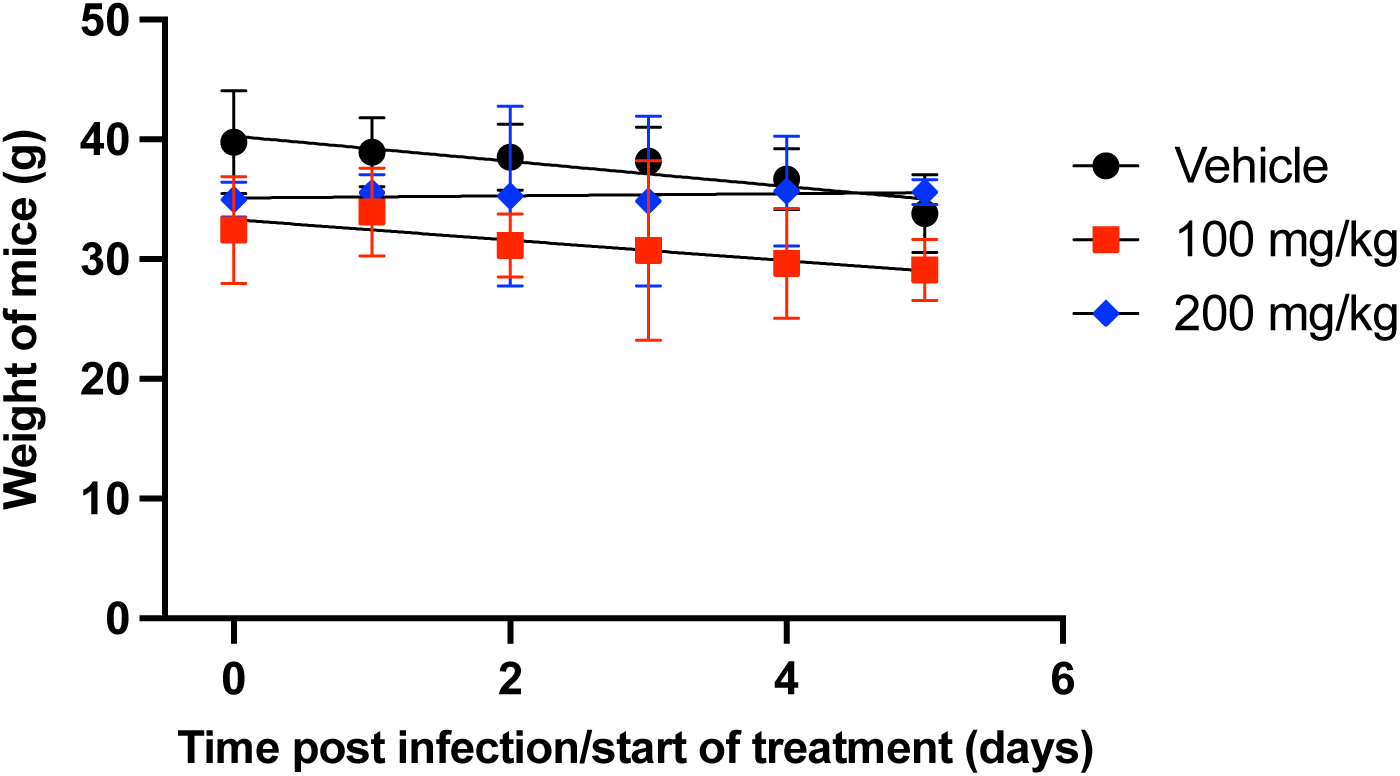
The weight of infected mice from the day of the start of treatment up to the day after the treatment period was completed. Mice were administered intraperitoneally with vehicle only (PBS; black circles), 100 mg/kg N3PT 9 (red squares), or 200 mg/kg N3PT 9 (blue diamonds), once a day for four days. Values are averaged from six mice. Error bars represent SD.

## Notes

### Competing Interest Statement

The authors have declared no competing interest.

